# Activity-dependent endoplasmic reticulum Ca^2+^ uptake depends on Kv2.1-mediated endoplasmic reticulum/plasma membrane junctions to promote synaptic transmission

**DOI:** 10.1101/2021.10.23.465516

**Authors:** Lauren C. Panzera, Ben Johnson, In Ha Cho, Michael M. Tamkun, Michael B. Hoppa

## Abstract

The endoplasmic reticulum (ER) forms a continuous and dynamic network throughout a neuron, extending from dendrites to axon terminals, and axonal ER dysfunction is implicated in several neurological disorders. In addition, tight junctions between the ER and plasma membrane (PM) are formed by several molecules including Kv2 channels, but the cellular functions of many ER-PM junctions remain unknown. Dynamic Ca^2+^ uptake into the ER during electrical activity plays an essential role in synaptic transmission as failure to allow rapid ER Ca^2+^ filling during stimulation activates stromal interaction molecule 1 (STIM1) and decreases both presynaptic Ca^2+^ influx and synaptic vesicle exocytosis. Our experiments demonstrate that Kv2.1 channels are necessary for enabling ER Ca^2+^ uptake during electrical activity as genetic depletion of Kv2.1 rendered both the somatic and axonal ER unable to accumulate Ca^2+^ during electrical stimulation. Moreover, our experiments show that the loss of Kv2.1 in the axon impairs synaptic vesicle fusion during stimulation via a mechanism unrelated to modulation of membrane voltage. Thus, our data demonstrate that the non-conducting role of Kv2.1 in forming stable junctions between the ER and PM via ER VAMP-associated protein (VAP) binding couples ER Ca^2+^ uptake with electrical activity. Our results further suggest that Kv2.1 has a critical function in neuronal cell biology for Ca^2+^-handling independent of voltage and reveals a novel and critical pathway for maintaining ER lumen Ca^2+^ levels and efficient neurotransmitter release. Taken together these findings reveal an essential non-classical role for both Kv2.1 and the ER-PM junctions in synaptic transmission.

**Significance:** The endoplasmic reticulum (ER) extends throughout the neuron as a continuous organelle, and its dysfunction is associated with several neurological disorders. During electrical activity, the ER takes up Ca^2+^ from the cytosol which has been shown to support synaptic transmission. This close choreography of ER Ca^2+^ uptake with electrical activity suggests functional coupling of the ER to sources of voltage-gated Ca^2+^ entry through an unknown mechanism. Here we report a non-conducting role for Kv2.1 through its ER binding domain that is necessary for ER Ca^2+^ uptake during neuronal activity. Loss of Kv2.1 profoundly disables neurotransmitter release without altering presynaptic voltage suggesting that Kv2.1-mediated signaling hubs play an important neurobiological role in Ca^2+^ handling and synaptic transmission independent of ion conduction.

## Introduction

The members of the Kv2 family of voltage-gated K^+^ (Kv) channels, Kv2.1 and Kv2.2 are widely expressed in neurons within the mammalian brain, with Kv2.1 dominating in hippocampal neurons (1-3). These channels play an important classical role repolarizing somatic membrane potential during high-frequency stimulation (4). However, Kv2 channels also form micron-sized clusters on the cell membrane, where they are largely non-conductive (5). When clustered, these non-conductive channels act as molecular hubs directing protein insertion and localization including during the fusion of dense-core vesicles (6-9), and as sites for the enrichment of voltage-gated Ca^2+^ channels (10). The Kv2 clustering mechanism is due to the formation of stable tethers between the cortical endoplasmic reticulum (ER) and the plasma membrane (PM) through a non-canonical FFAT motif located on the Kv2 C-terminus, which interacts with VAMP-associated protein (VAP) embedded in the ER membrane (11). These Kv2.1-mediated junctions between the ER and PM are in close (∼15 nm) proximity (12), forming critical Ca^2+^-signaling domains that have been conserved from yeast to mammals (12-14) and are necessary to cluster Kv2.1 channels. ER-PM junctions are formed by many types of proteins, although most are ER proteins that transiently interact with specific lipids on the PM (reviewed previously (15)). The purpose of the Kv2.1-VAP mediated ER-PM junctions is not functionally understood in neurons to date.

Cytosolic Ca^2+^ is essential for initiating multiple cell functions including secretion, muscle contraction, proliferation, apoptosis, and gene expression (reviewed previously (16)). However, Ca^2+^ is also strongly buffered, especially in most neurons, and often requires local Ca^2+^ exchange between channels and pumps localized to organelles and the PM. The ER plays a central role in both Ca^2+^ signaling and storage, and dysfunction of ER morphology and Ca^2+^ handling has been linked to several unique neurological pathologies including hereditary spastic paraplegia (17), Alzheimer’s disease (18), and amyotrophic lateral sclerosis (19). Currently, the only known cellular mechanism used to replenish ER Ca^2+^ stores is through activation of store-operated Ca^2+^ entry (SOCE). Depletion of the ER’s lumenal Ca^2+^ is sensed by stromal interaction molecule 1 (STIM1), which aggregates and concentrates Orai proteins on the PM to initiate Ca^2+^ influx through Ca^2+^ release-activated Ca^2+^ (CRAC) channels. Recent studies, however, have revealed that a second frequently accessed pathway exists in neurons where stimulation-evoked Ca^2+^ influx is rapidly taken up by the ER during neuronal activity, rather than in reaction to severe depletion of ER lumen Ca^2+^. Failure to quickly increase lumenal Ca^2+^ during action potential (AP) firing leads to ER Ca^2+^ depletion and impaired synaptic vesicle fusion (20). Thus, lumenal ER Ca^2+^ plays an essential role in maintaining synaptic transmission in active healthy neurons suggesting that a mechanism other than SOCE must be important for neuronal communication.

Taken together, Kv2.1 clusters have been shown to localize L-type voltage-gated calcium channels at the PM while also anchoring the ER in close proximity to the PM (10). We hypothesized that these Kv2.1-mediated ER-PM junctions are uniquely positioned to serve a critical role as dynamic signaling domains for rapid ER Ca^2+^ uptake during electrical activity in neurons. We measured Kv2.1’s role in ER Ca^2+^-handling using ER-GCaMP6-150 and found that AP-evoked Ca^2+^ entry into the somatic ER was absent with genetic depletion of Kv2.1 channels. This non-conducting role of Kv2.1 which enables ER-Ca^2+^ filling also requires sarco/endoplasmic reticulum Ca^2+^-ATPase (SERCA) pumps. Moreover, we demonstrate a novel non-conducting role for Kv2.1 in the axon that is essential for enabling ER-Ca^2+^ uptake during electrical activity. We go on to show that depletion of Kv2.1 impaired overall synaptic physiology through decreased presynaptic Ca^2+^ entry and synaptic vesicle exocytosis. Finally, we demonstrate that this role requires Kv2.1’s C-terminal VAP-binding domain to restore synaptic transmission.

## Results

### Kv2.1 channels have both conducting and non-conducting roles in the soma

Kv2.1 channels are widely expressed in excitatory and inhibitory neurons in both the hippocampus (21-24) and cortex (24-27), where they have two prominent functions: One, a conducting role repolarizing somatic membrane potential during AP firing (3, 4); and two, a scaffolding role forming ER-PM junctions with the ER-resident protein VAP (11). We sought to quantitatively address Kv2.1 function in both roles. To measure a role in somatic AP repolarization, we first used a combination of shRNA targeting the endogenous Kv2.1 channel and the optical voltage indicator QuasAr (28, 29) in cultured hippocampal neurons. Genetic knockdown of Kv2.1 by shRNA resulted in a 76% reduction in immunostained Kv2.1 fluorescence intensity (Fig. S1A and S1B). We also expressed the shRNA using adeno-associated virus (AAV) for high efficiency transduction and found an 80% reduction in Kv2.1 protein expression (Fig. S1C). Consistent with Kv2.1’s role as a delayed rectifying potassium channel we observed no change in the AP amplitude, but an increase (14.5%) in the full-width at half maximum (FWHM) of APs recorded in the soma of cultured hippocampal neurons lacking the Kv2.1 channel (Control neurons, 2.01 ± 0.097 ms; Kv2.1 KD neurons, 2.30 ± 0.093 ms; *p*<0.05) (Fig. 1A-C).

**Figure 1.**
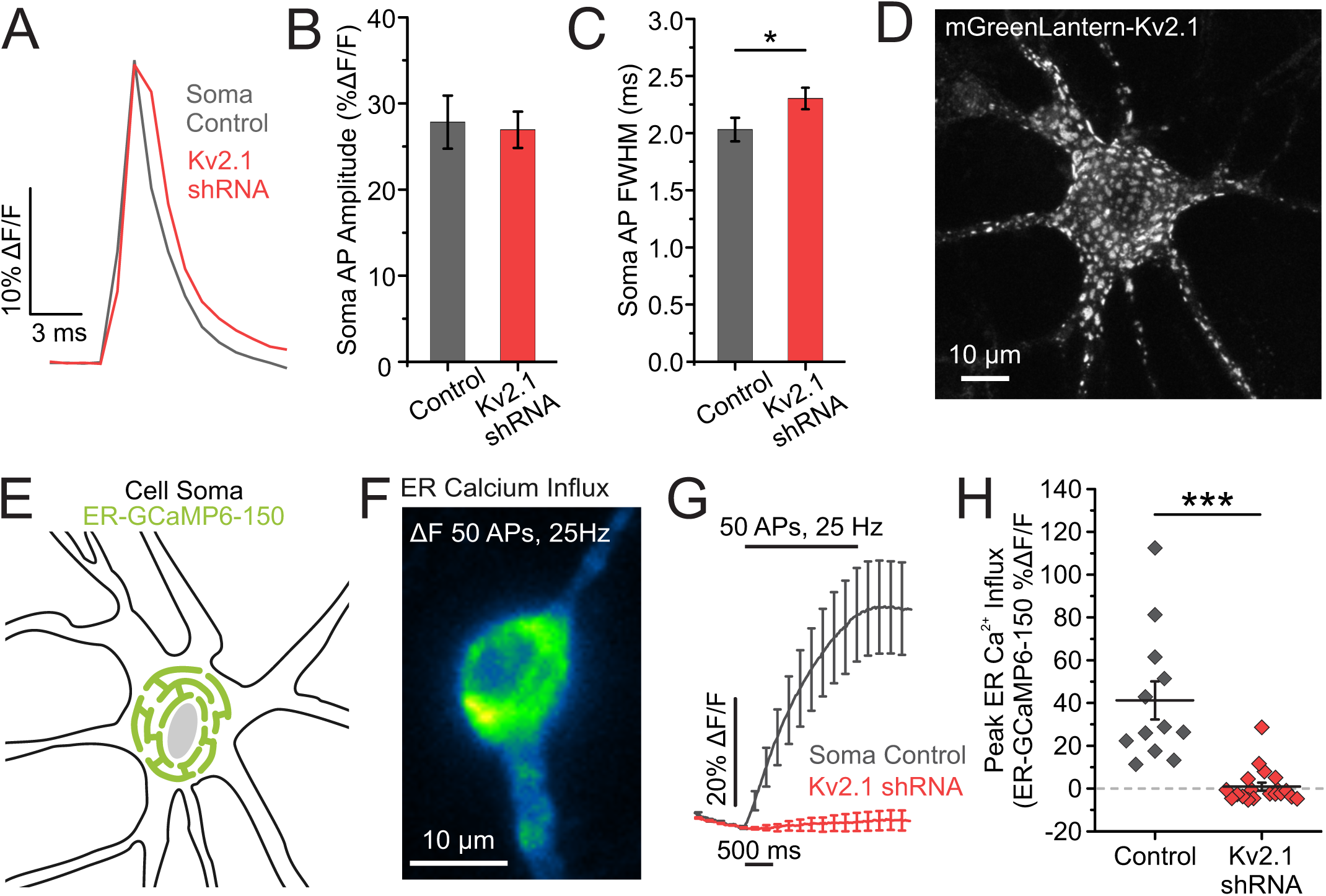
Kv2.1 has both ionotropic and non-ionotropic functions in the soma. **(A)** Average traces of somatic QuasAr fluorescence, trial averaged from 100AP stimulations. **(B-C)** Quantification of AP amplitude **(B)** and full width at half maximum (FWHM) **(C)** (Control neurons, *n* = 11 cells; Kv2.1 KD neurons, *n* = 10 cells; *p*<0.05 for FWHM comparison, Student’s *t*-test). **(D)** Example image of a cultured hippocampal neuron expressing mGreenLantern-Kv2.1. Note distinct clusters form across the membrane surface. **(E)** Cartoon of a neuronal soma expressing the fluorescent calcium indicator ER-GCaMP6-150 in the ER lumen. **(F)** Image of the change in fluorescence of somatic ER-GCaMP6-150 in response to a train of stimulation. **(G-H)** Average fluorescence traces of somatic ER-GCaMP6-150 **(G)** and quantification of peak fluorescence **(H)** for both control and Kv2.1 knockdown neurons (Control neurons, *n* = 12 cells; Kv2.1 KD neurons, *n* = 19 cells; *p*<0.001, Student’s *t*-test).

While conductive Kv2.1 channels have homogenous membrane expression, endogenous and transfected Kv2.1 channels also prominently localize in micron-sized clusters within the somatodendritic compartments and axon initial segment both *in vivo* and *in vitro* (30-32), and have been thought to be excluded from the distal axon and presynaptic terminals (24). An example is seen in a hippocampal neuron expressing Kv2.1 tagged with mGreenLantern (33), where clustering is obvious in the soma, proximal dendrites and axon initial segments of our cultured hippocampal neurons, consistent with previous findings (24, 34, 35) (Fig. 1D). Previously, we and others demonstrated that Kv2.1 localization is necessary and sufficient to induce ER-PM junctions across cell types, including HEK cells and developing hippocampal neurons (12, 36). Further, TIRF imaging in young cultured hippocampal neurons showed retraction of cortical ER following declustering of Kv2.1 channels (12), indicating that both the expression and placement of Kv2.1 within the membrane is important for junction formation.

Given the role for Kv2.1 clusters in both localizing voltage-gated Ca^2+^ channels (10) and forming ER-PM junctions (11) we were curious about the effect of reducing Kv2.1 expression on ER Ca^2+^ uptake during electrical activity. We expressed the low-affinity ER Ca^2+^ indicator ER-GCaMP6-150 (ER-GCaMP) in cultured hippocampal neurons and measured ER Ca^2+^ influx in the soma of transfected neurons during stimulation (Fig. 1E-F). While a train of 50 APs normally causes a robust increase in ER Ca^2+^, this process was severely impaired (97.64%) by the loss of Kv2.1 (Control neurons, 41.21 ± 8.89% ΔF/F; Kv2.1 KD neurons, 0.97 ± 1.87% ΔF/F; *p*<0.001) (Fig. 1G-H). To ensure that this uptake was mediated by SERCA pumps we applied the SERCA inhibitor cyclopiazonic acid (CPA), which completely blocked ER Ca^2+^ filling (Control neurons, 40.85 ± 7.42% ΔF/F; CPA-treated control neurons, -7.74 ± 0.84% ΔF/F; *p*<0.01) (Fig. S2A, C). Importantly, ER Ca^2+^ influx in Kv2.1 knockdown neurons was impaired to the extent that SERCA inhibition had little effect (Kv2.1 knockdown neurons, 2.21 ± 3.51% ΔF/F, CPA-treated knockdown neurons, -3.52 ± 1.34% ΔF/F) (Fig. S2B-C). We also examined the effect of Kv2.1 on somatic ER Ca^2+^ refilling following store depletion with Ca^2+^-free external solutions in developing neurons (8 DIV) before neurons express significant levels of Kv2.1 channels (30, 37). TIRF imaging was used to optically isolate ER/PM junctions and ER Ca^2+^ levels were measured with the ER-targeted fluorescent Ca^2+^ indicator CEPIA*er* (38) with and without Kv2.1 transfection as illustrated in Fig. S3A. After reintroducing extracellular Ca^2+^ the refilling rate of the cortical ER in Kv2.1-expressing neurons was 5-fold greater than that observed in neurons without Kv2.1 (Fig. S3B). To test whether ER-PM junction formation through VAP recruitment was sufficient to increase the rate of ER Ca^2+^ refilling, we used a chimeric approach. VAPs are typically diffusely localized in the ER and expressing the Kv2.1 C-terminal tail including the non-canonical FFAT VAP-binding domain fused to a single-pass transmembrane glycoprotein (CD4) is sufficient to redistribute VAPs near the plasma membrane of HEK cells and neurons (11). The refilling rate at ER/PM contact sites formed by the CD4-Kv2.1FFAT motif chimera was half that of Kv2.1-induced ER/PM junctions (Fig. S3B). This intermediate value suggests that simply forming ER-PM junctions does not confer efficient ER-Ca^2+^ uptake, and Kv2.1 functionalizes the ER-PM junction with regards to Ca^2+^ handling.

Taken together, our results confirm the canonical ionotropic role of Kv2.1 channels at the soma for regulating membrane voltage, while also revealing a novel non-conducting role for Kv2.1 in regulating ER Ca^2+^ filling during electrical activity or following ER store depletion.

### Endogenous and transfected Kv2.1 localize beyond the somatodendritic compartment into axons and presynaptic compartments

Somatic Kv2.1 clearly regulates ER Ca^2+^ stores as illustrated in Figs. 1, S2 and S3. Since ER Ca^2+^ regulates axonal glutamate release we wondered whether Kv2 channels could influence Ca^2+^ homeostasis, and thus glutamate release, in this neuronal compartment. However, the Kv2 channel dogma states that these channels are only found on the neuronal soma, axon initial segment, and within proximal dendrites (30, 31, 39). Therefore, we first sought to determine whether Kv2.1 channels could be found localized over the ER at presynaptic sites.

The first set of experiments sought to immunolocalize endogenous Kv2.1 and its similar family member, Kv2.2, within axons of our neuronal cultures. As illustrated in Fig. 2A, Kv2.1 can be detected in synapsin-positive presynaptic compartments in DIV 16 hippocampal cultures (see white arrows in Fig. 2A inset). The level of expression is much lower than that observed on the soma and, if detected by investigators previously, was probably viewed as non-specific antibody staining. As demonstrated in Fig. S4A and B, use of a dominant-negative construct which blocks Kv2.1 trafficking to the neuronal surface prevented detection of Kv2.1 on both the somatic surface and in axonal compartments, validating the axonal immunolabeling. Kv2.2 immunostaining was also detected weakly in both somatic and axonal compartments (Fig. S5A-B) and this was also blocked following expression of the dominant-negative construct (see Fig. S5C and D). Note that while some Kv2.2 immunostaining was present in presynaptic compartments (see white arrows in Fig. S5A inset) it often appeared at a much lower level compared to Kv2.1, and Kv2.2 was at times not detected. In addition, Kv2.2 immunolabeling was often found adjacent to the synapsin puncta as opposed to fully co-localizing (see yellow arrowheads in Fig. S5A inset). Perhaps Kv2.2 predominates in postsynaptic (dendritic) compartments while present at a much lower level relative to Kv2.1 on the presynaptic side. However, given that the anti-Kv2.2 antibody is likely less efficient than its Kv2.1 counterpart, this issue remains an open question at this time. Still, these immunolabeling data indicate both Kv2.1 and 2.2 are expressed in both somatic and presynaptic compartments in our neuronal cultures and surface trafficking of both variants is impaired by expression of the DN form of Kv2.1.

**Figure 2.**
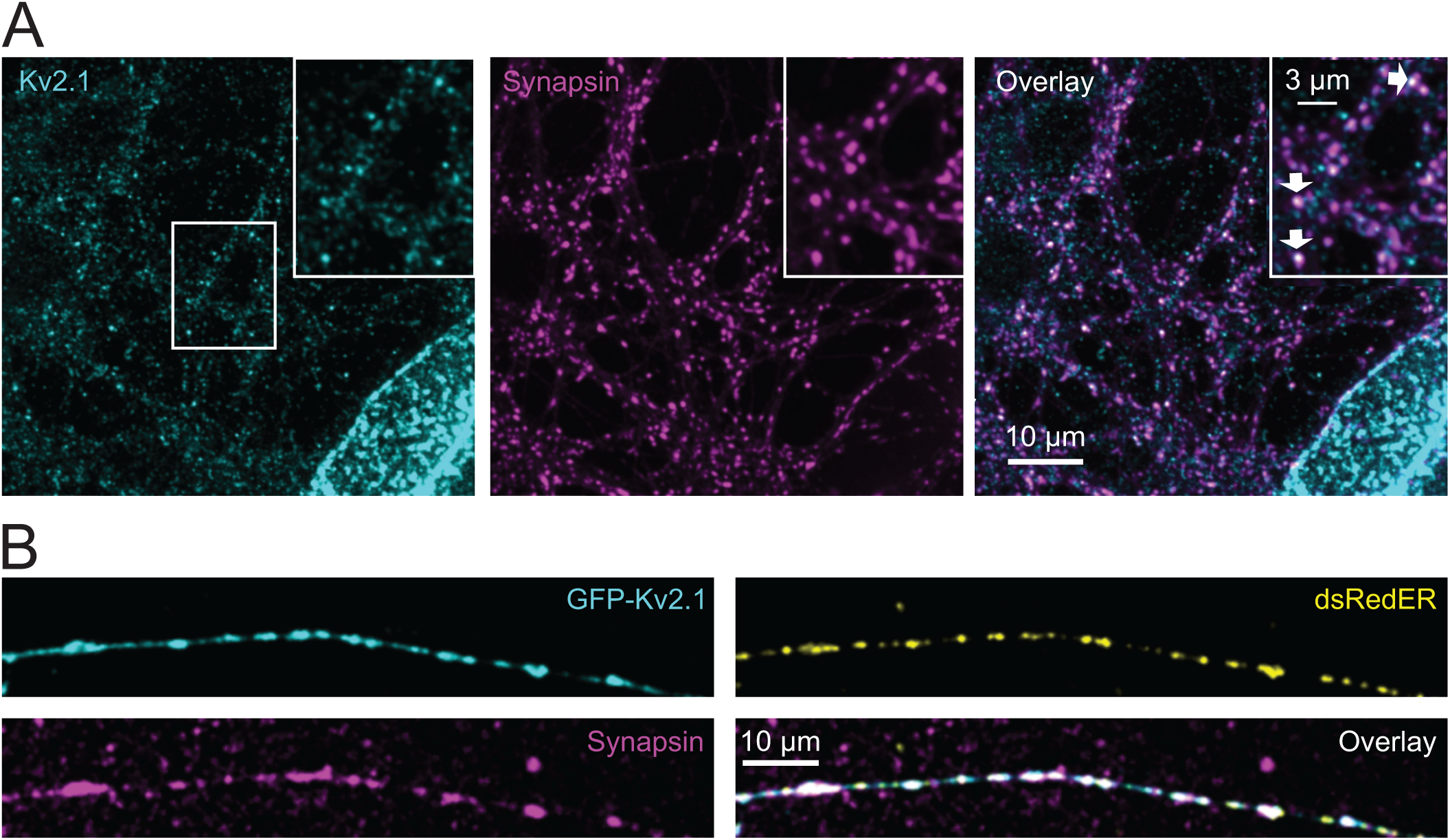
Endogenous Kv2.1 localizes beyond the somatodendritic compartment into axons and terminals. **(A)** Immunolabeled images of endogenous Kv2.1 (*cyan*) and synapsin (*magenta*), with merged channels (*right*), in DIV 14 neurons. The center white box indicates the region enlarged as shown in the inset. Arrows indicate Kv2.1 colocalized with synapsin-positive presynaptic terminals. **(B)** Images of transfected and fixed DIV 16 neurons expressing GFP-Kv2.1 (*cyan*) and dsRedER (*yellow*), with immunolabeled synapsin (*magenta*).

We next transfected neurons with GFP-Kv2.1 and dsRedER followed by fixation and immunolabeling for synapsin. As shown in Fig. 2B, the transfected Kv2.1 localized to presynaptic compartments that were enriched in the ER luminal marker dsRedER. In summary, we were surprised to discover a population of Kv2 channels that existed beyond the somatodendritic and axon initial segment compartments, with both the endogenous Kv2 channels and the transfected Kv2.1 clearly present in presynaptic compartments. This localization is not fully unexpected, however, given that ER/PM junctions have been reported in axon terminals (40) and the only known localization mechanism for Kv2 channels involves tethering to ER VAPs.

### Loss of Kv2.1 impairs axonal ER Ca^2+^ influx during stimulation independent of ion conduction

We next investigated the functional role of axonal Kv2.1 channels. During our previous efforts to understand modulation of the AP waveform in cultured hippocampal neurons, we found that presynaptic terminals primarily rely on Kv1 and Kv3 channels to repolarize the AP (41). Consistent with this finding, optical recordings of axonal AP waveforms uncovered no difference in the amplitude or FWHM between control and Kv2.1 KD neurons (Fig. 3A-C). Next, we confirmed previous findings that the axonal ER takes up Ca^2+^ during AP stimulation and again observed a robust increase in lumenal Ca^2+^ from SERCA pump activation during trains of stimulation (Fig. 3D-G). This process of axonal ER Ca^2+^ filling during neuronal activity also appeared to rely on Kv2.1 channels as neurons transfected with Kv2.1 shRNA were severely impaired in ER Ca^2+^ uptake during trains of stimulation identical to the soma (Control neurons, 33.04 ± 7.63% ΔF/F; Kv2.1 KD neurons, 5.26 ± 2.97% ΔF/F; *p*<0.01) (Fig. 3F-G).

**Figure 3.**
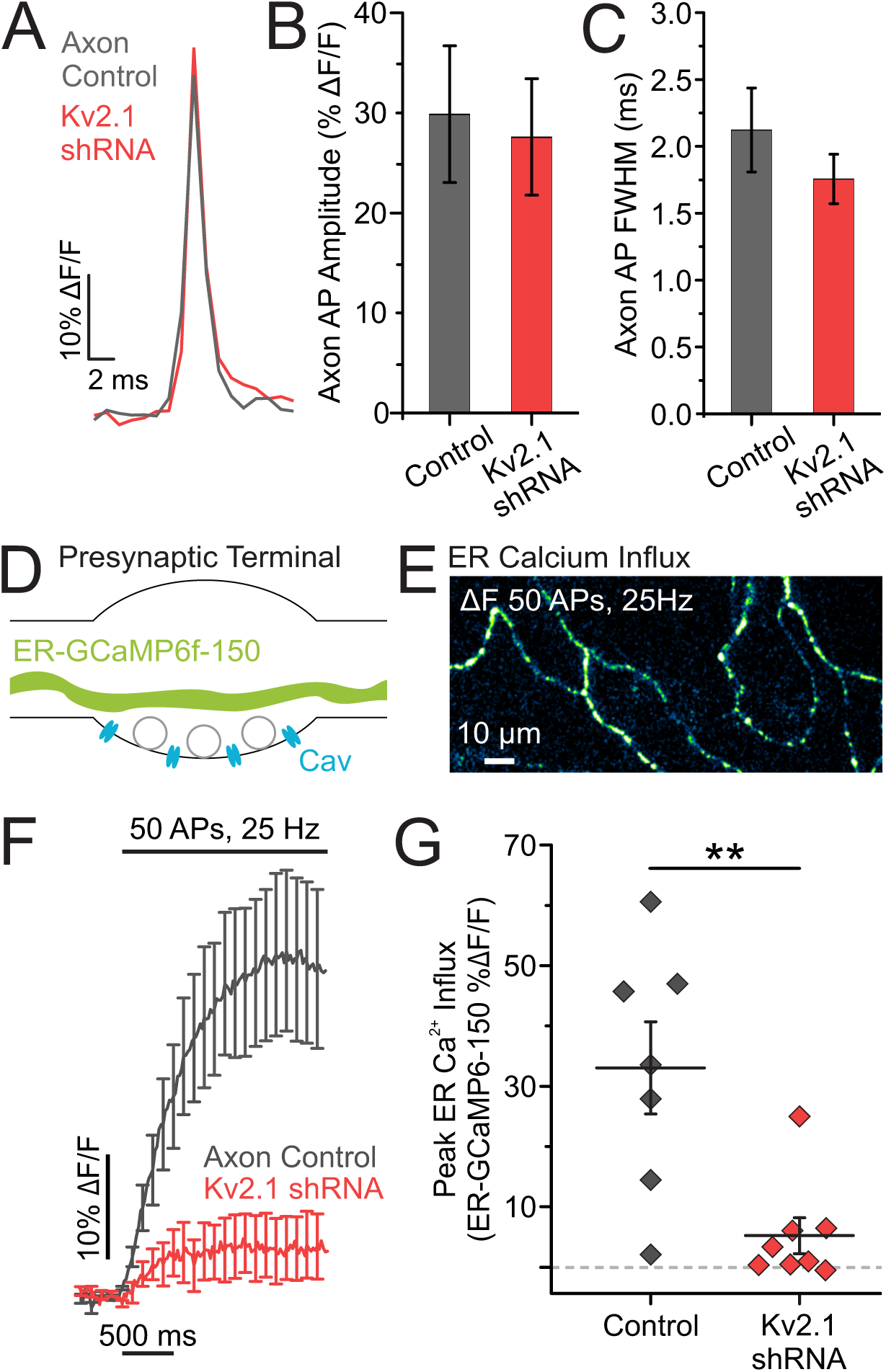
Loss of Kv2.1 impairs axonal ER calcium influx during stimulation. **(A)** Representative traces of axonal QuasAr fluorescence, trial averaged from 100 AP stimulations. **(B-C)** Quantification of AP amplitude **(B)** and full width at half maximum **(C)** (Control neurons, *n* = 4 cells; Kv2.1 KD neurons *n* = 6 cells). **(D)** Cartoon of a presynaptic terminal expressing the fluorescent calcium indicator ER-GCaMP6-150 in the ER lumen. **(E)** Image of the change in fluorescence of axonal ER-GCaMP6-150 in response to a train of stimulation. **(F-G)** Average fluorescence traces of axonal ER-GCaMP6-150 **(F)** and quantification of peak fluorescence **(G)** for both control and Kv2.1 knockdown neurons (Control neurons, *n* = 7 cells; Kv2.1 KD neurons, *n* = 8 cells; *p*<0.01, Student’s *t*-test).

Since both Kv2.1 and Kv2.2 localized to presynaptic compartments where they form ER-PM junctions, we wanted to determine the level of contribution from total Kv2 channels to regulate axonal ER Ca^2+^. We used the dominant-negative construct (Kv2.1 DN) illustrated in Fig. S4 that prevents both Kv2.1 and Kv2.2 from being expressed at the PM (42). We found that depletion of surface Kv2.1 and Kv2.2 channels dramatically decreased axonal AP-evoked ER Ca^2+^ influx (Control neurons, 18.17 ± 4.99% ΔF/F; Kv2.1 DN neurons, 3.18 ± 2.17% ΔF/F; *p*<0.05) (Fig. S6A-B). Importantly, the additional loss of Kv2.2 did not substantially change the percent decrease in ER-GCaMP responses (Kv2.1 shRNA KD neurons 84.08% decrease; Kv2.1 DN neurons, 82.50% decrease), indicating Kv2.1 plays the dominant role in axonal ER Ca^2+^ influx in our preparation as implied by the weak presynaptic immunolabeling of Kv2.2 relative to that observed with Kv2.1. Together, these results suggest that while Kv2.1 does not play an ionotropic role in the axon, it does play a novel and critical role with regards to ER-Ca^2+^ handling.

### Presynaptic Kv2.1 modulates neurotransmission independently of conduction

This striking non-ionotropic role for axonal Kv2.1 channels in regulating ER Ca^2+^ stores was especially intriguing in light of recent work demonstrating that blocking SERCA pumps or decreasing ER Ca^2+^ stores impairs neurotransmission (20). To explore a non-conducting role for Kv2.1 in modulating vesicle fusion, we used the pH-sensitive reporter of synaptic vesicle exocytosis (pHluorin) fused to the vesicular glutamate transporter (vGlut-pHluorin). vGlut-pHluorin signals are reported as a percentage of the total vesicle pool (% exocytosis), whose fluorescence is obtained by perfusion of a Tyrode’s solution containing 50mM NH_4_Cl buffered at pH 7.4 using 25mM HEPES (Fig. 4A-B) (43-46). Neurons transfected with Kv2.1 shRNA had a large (43.6%, *p*<0.0001) reduction in exocytosis (Fig. 4C-D), consistent with impaired Ca^2+^ uptake into the ER. These results suggest that Kv2.1 plays a significant non-conducting role in neurotransmission.

**Figure 4.**
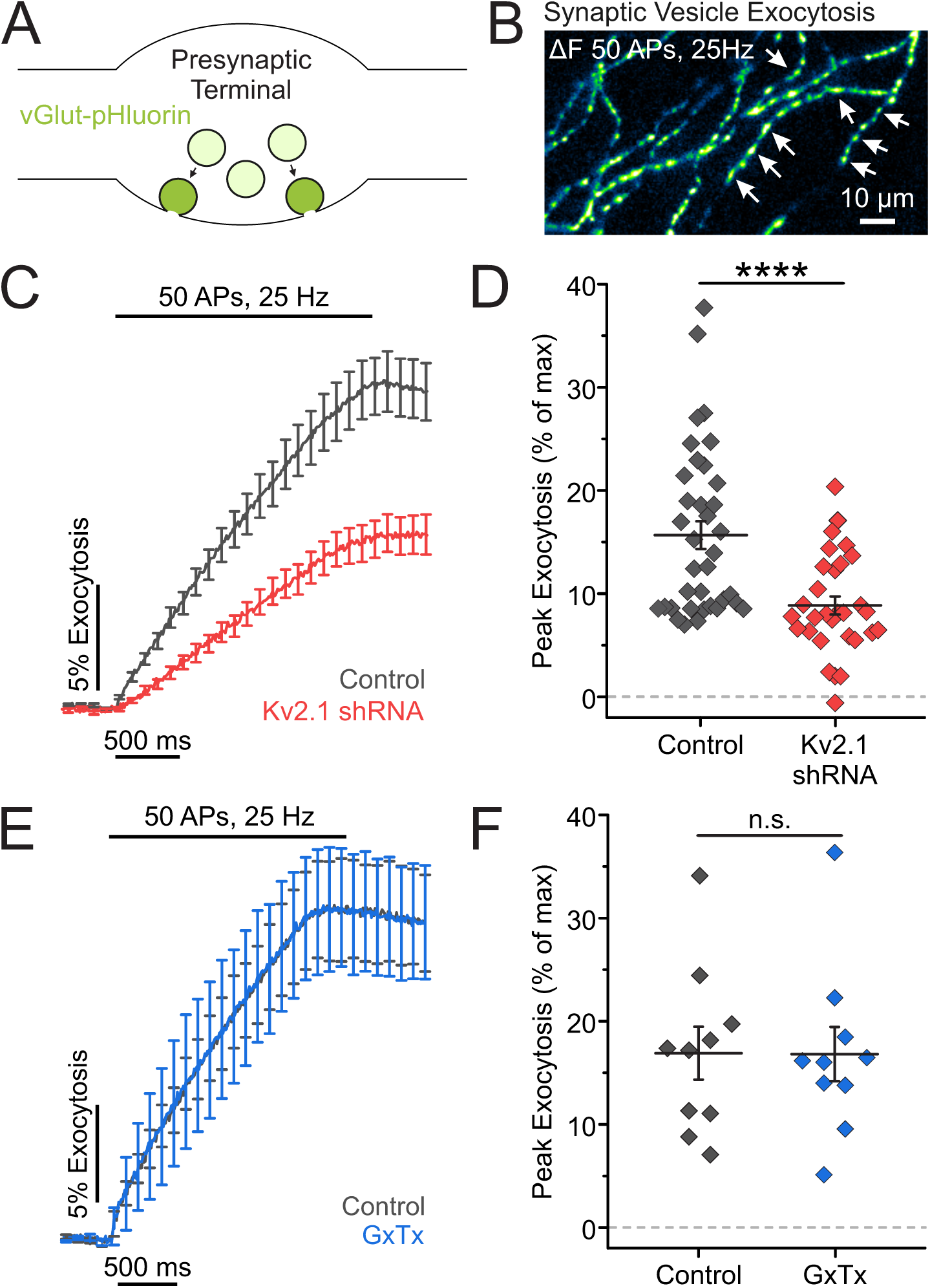
Presynaptic Kv2.1 modulates neurotransmission independently of conduction. **(A)** Cartoon of a presynaptic terminal containing synaptic vesicles expressing vGlut-pHluorin, a pH-sensitive indicator of exocytosis. **(B)** Image of the change in fluorescence of vGlut-pHluorin in response to a train of stimulation. Arrows mark locations of presumptive presynaptic terminals. **(C-D)** Average fluorescence traces of vGlut-pHluorin **(C)** and quantification of peak fluorescence **(D)** for both control and Kv2.1 knockdown neurons (Control neurons, *n* = 36 cells; Kv2.1 KD neurons, *n* = 30 cells; *p*<0.0001, Student’s *t*-test). **(E-F)** Average fluorescence traces of vGlut-pHluorin **(E)** and quantification of peak fluorescence **(F)** for both control and Guangxitoxin-1 E (GxTx) treated neurons (*n* = 10 cells, paired *t*-test).

To confirm that this result was due to the loss of Kv2.1 protein rather than loss of a conducting, ionotropic or voltage-sensing role, we turned to pharmacology. Kv2 channels detect membrane potential changes through a group of positive charges located in the S4 domain of the channel α subunit. We used the gating modifier Guangxitoxin-1E (GxTx) to block Kv2.1 voltage-sensing and conduction. GxTx induces a depolarizing shift in the voltage-dependent activation of Kv2.1 with high potency and selectivity [IC_50_: 0.71 nM, (47)]. Perfusion of 100nM GxTx to prevent potassium conduction through Kv2.1 did not affect vGlut-pHluorin responses (Fig. 4E-F). Together with the finding that loss of Kv2.1 had no effect on axonal membrane voltage during an AP, these results further demonstrate that axonal Kv2.1 modulates neurotransmission independently of potassium conduction.

### Reducing Kv2.1 expression impairs presynaptic Ca^2+^ influx

It was previously shown that chronically blocking SERCA pumps and ER Ca^2+^ uptake drives a STIM1-based feedback loop that inhibits Ca^2+^ influx from voltage-gated Ca^2+^ channels and vesicle fusion (20). We sought to determine if a similar pathway was engaged with the loss of Kv2.1-based ER Ca^2+^ refilling during electrical activity as a measure of potential ER distress from the loss of Kv2.1. We measured how depletion of Kv2.1 altered presynaptic Ca^2+^ influx during trains of stimulation using the fluorescent Ca^2+^ indicator GCaMP6f fused to synaptophysin (48, 49) (SypGCaMP6f) (Fig. 5A-B). Not surprisingly, compared to controls, neurons co-transfected with SypGCaMP6f and Kv2.1 shRNA had reduced presynaptic Ca^2+^ influx when stimulated with 50 APs delivered at 25 Hz (Control neurons, 288.60 ± 36.06% ΔF/F; Kv2.1 KD neurons 186.84 ± 18.99% ΔF/F; *p*<0.05) (Fig. 5C-D). Thus, the loss of Kv2.1 phenocopies the effects of blocking SERCA pumps with CPA. These results suggest that disabling ER Ca^2+^ uptake by the loss of Kv2.1 during electrical activity feeds back into general Ca^2+^ homeostasis and synaptic transmission.

**Figure 5.**
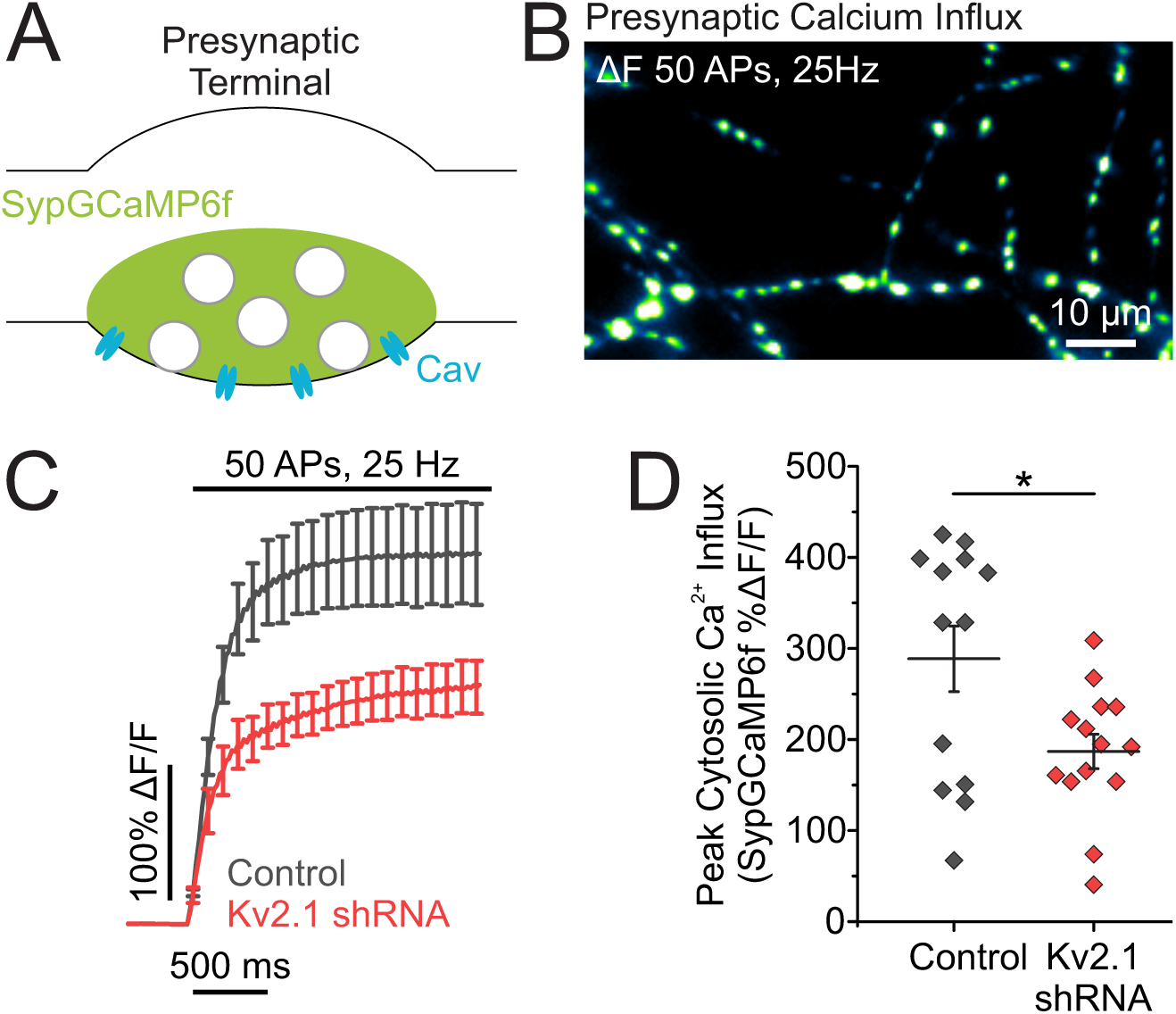
Reducing Kv2.1 expression impairs evoked presynaptic calcium influx. **(A)** Cartoon of a presynaptic terminal expressing the fluorescent calcium indicator Synaptophysin-GCaMP6f (SypGCaMP6f). **(B)** Image of the change in fluorescence of SypGCaMP6f in response to a train of stimulation. **(C-D)** Average fluorescence traces of SypGCaMP6f **(C)** and quantification of peak fluorescence **(D)** in both control and Kv2.1 knockdown neurons (Control neurons, *n* = 13 cells; Kv2.1 KD neurons, *n* = 14 cells; *p*<0.05, Student’s *t*-test).

### The Kv2.1 channel C-terminus is necessary for maintaining synaptic transmission

We next addressed the mechanism underlying Kv2.1’s role in ER Ca^2+^ handling and neurotransmission by conducting rescue experiments. To validate our rescue approach, we introduced three silent point mutations into the coding sequence of Kv2.1 at the shRNA target site. Expression of this “wobbled” Kv2.1 (wKv2.1) in the Kv2.1 KD background was sufficient to fully rescue the synaptic vesicle exocytosis defect, confirming the specificity of our shRNA (Kv2.1 KD neurons, 6.29 ± 0.89%; wKv2.1 rescue neurons, 12.22 ± 2.40% exocytosis; *p*<0.05) (Fig. 6A-C).

**Figure 6.**
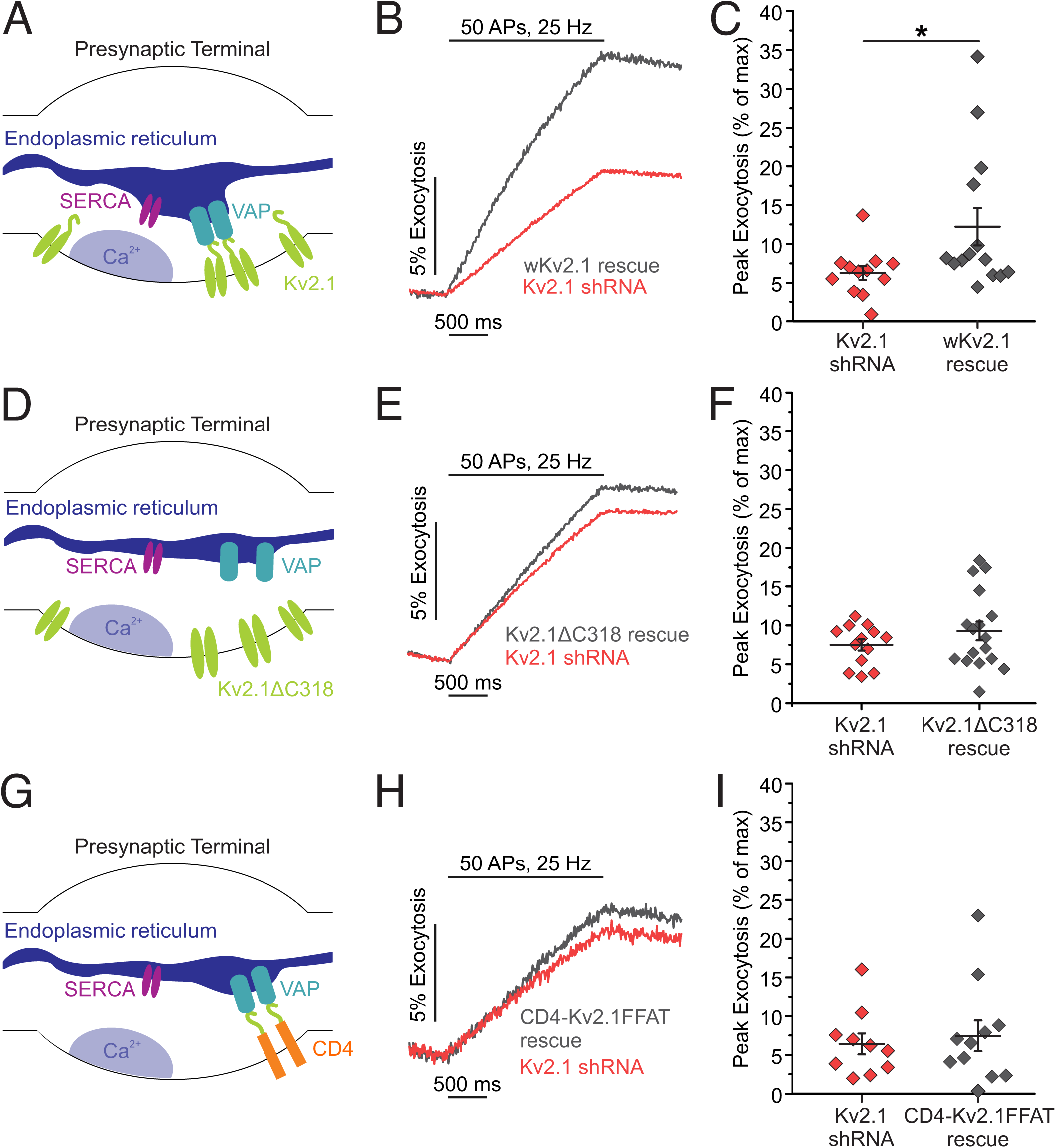
The VAP-binding domain of Kv2.1 is necessary to rescue synaptic vesicle exocytosis. **(A)** Cartoon of a presynaptic terminal with endogenous Kv2.1 channels tethering the ER to the plasma membrane, positioning SERCA channels nearby to a source of presynaptic calcium influx. **(B-C)** Average fluorescence traces of vGlut-pHluorin **(B)** and quantification of peak fluorescence **(C)** in both Kv2.1 and shRNA-resistant (wobbled) Kv2.1 rescue neurons (Kv2.1 KD neurons, *n* = 12 cells; wKv2.1 rescue neurons, *n* = 14 cells; *p*<0.05, Student’s *t*-test). **(D)** Cartoon of a presynaptic terminal expressing mCherry-Kv2ΔC318, which is missing the Kv2.1 VAP-binding domain and does not localize SERCA near sites of presynaptic calcium influx. **(E-F)** Average fluorescence traces of vGlut-pHluorin **(E)** and quantification of peak fluorescence **(F)** in both Kv2.1 shRNA and Kv2ΔC318 rescue neurons (Kv2.1 KD neurons, *n* = 13 cells; Kv2ΔC318 neurons, *n* = 17 cells). **(G)** Cartoon of a presynaptic terminal expressing the CD4-Kv2.1FFAT chimera, which creates ER-plasma membrane junctions perhaps in alternate locations away from sites of presynaptic calcium influx. **(H-I)** Average fluorescence traces of vGlut-pHluorin **(H)** and quantification of peak fluorescence **(I)** in both Kv2.1 shRNA and CD4-Kv2.1FFAT rescue neurons (Kv2.1 KD neurons, *n* = 10 cells; CD4-Kv2.1FFAT neurons, *n*= 11 cells).

The formation of ER-PM junctions by Kv2.1 channels in the PM occurs via a noncanonical VAP-binding motif within the C-terminal tail of Kv2.1 (11, 36); deletion or mutation of this motif abolishes both Kv2.1 clusters and ER-PM junctions (34, 39). However, despite a lack of clustering, the loss of the C-terminal tail does not alter Kv2.1’s electrical function (6, 50). Not surprisingly, without the C-terminal tail that enables VAP binding and Kv2.1 channel clustering, we could no longer visualize punctate Kv2.1 structures in the soma or axon. Moreover, expressing Kv2.1 with a truncated C-terminus (Kv2ΔC318) in the Kv2.1 KD background was also unable to restore synaptic transmission (Kv2.1 KD neurons, 7.49 ± 0.72%; Kv2ΔC318 neurons, 9.28 ± 1.21% exocytosis) (Fig. 6D-F). Next, we were curious if the C-terminus alone is sufficient to restore synaptic function. To distinguish between the specialized Kv2.1 ER-PM junctions and simply bringing ER and PM membranes together through the VAP binding domain, we returned to the CD4-Kv2.1FFAT chimera used in Fig. S3. We transfected this fusion protein consisting of the C-terminus of Kv2.1 including the VAP-binding domain appended to the transmembrane protein CD4 and tested whether the Kv2.1 VAP-binding domain alone was sufficient to rescue the synaptic vesicle phenotype (Fig. 6G). Expression of the CD4-Kv2.1FFAT was localized to the presynaptic terminals (Fig. S7), however it was not sufficient to restore exocytosis to control levels in Kv2.1 knockdown neurons (Kv2.1 KD neurons, 6.41 ± 1.35%; CD4-Kv2.1FFAT neurons, 7.45 ± 1.98% exocytosis) (Fig. 6H-I). This result indicates that other sequences within Kv2.1 are required to enable efficient ER Ca^2+^ uptake that can support synaptic transmission, rather than acting simply to recruit VAP. Further, this observation is in agreement with our finding that ER Ca^2+^ uptake after store depletion is much less efficient when expressing CD4-Kv2.1FFAT instead of full length Kv2.1 (Fig. S3). At the same time, Kv2.1 clustering and inclusion of the VAP-binding motif is essential, as expressing Kv2.1 with a truncated C-terminus (Kv2ΔC318) was also unable to restore synaptic transmission. Taken together, these results support the role of Kv2.1 as an essential hub protein that provides a novel mechanism to enable efficient ER Ca^2+^ uptake during electrical stimulation and has an important function in maintaining neurotransmitter release.

## Discussion

In most cells, the ER acts mainly as a Ca^2+^ source when different pathways activate ryanodine receptors or inositol 1,4,5-trisphosphate (IP3) receptors. Intriguingly, the neuronal ER acts as a net Ca^2+^ sink in both the soma and axon of neurons, using SERCA pumps to extract Ca^2+^ from small microdomains in the cytosol formed by the opening of voltage-gated Ca^2+^ channels. Inhibiting the activity of SERCA during electrical activity has previously been demonstrated to dramatically impair synaptic function (51, 52) due to what was recently identified as the perturbation of STIM1 proteins (20). To date, a mechanism that enables efficient access by SERCA to sources of PM voltage-gated Ca^2+^ influx has not been identified. Here, we provide evidence that the non-conducting signaling hubs of Kv2.1 channels enable this elegant coupling of Ca^2+^ uptake into the ER during electrical activity in both the soma and synaptic terminals (Figs. 1 and 3). Loss of Kv2.1 renders the ER unable to extract Ca^2+^ from the cytosol during electrical activity and makes the neuron behave as if the SERCA pumps are not functional during stimulation. Interestingly, the loss of Kv2.1 channels matches the phenotypes of the SERCA block with respect to impaired cytosolic Ca^2+^ influx and vesicle fusion during electrical stimulation (Figs. 4 and 5). Interestingly, impaired AP-evoked calcium influx was also reported by another group when they knocked down VAP protein using shRNA (53). Our results demonstrate the importance of regulating internal Ca^2+^ stores for maintaining neurotransmission and support the necessity of Kv2.1-VAP ER-PM junctions, as the loss of the C-terminal VAP binding domain was unable to restore synaptic transmission (Fig. 6). Although the C-terminus alone was shown to recruit VAP and form ER-PM junctions in neurons and heterologous cells, our CD4-Kv2.1FFAT construct containing the VAP-binding domain also could not restore synaptic transmission and poorly supported somatic ER Ca^2+^ refilling (Figs. 6 and S3). While resolution limitation in fluorescent microscopy precludes us from making precise measurements of subsynaptic localization differences between Kv2.1 and CD4-Kv2.1FFAT, our results suggest that Kv2.1 truly acts a hub either by localizing sites of Ca^2+^ entry or through other interactions to position ER-PM junctions specifically proximal to Ca^2+^ entry to promote synaptic transmission (Fig. 6). Taken together, these results support a non-conducting role for the Kv2.1 channel as a hub-protein to couple ER Ca^2+^ handling with electrical activity essential for neuronal function. Kv2.1 has also been identified to play a non-conducting role in exocytosis/secretion in other cells, especially those that are electrically active, e.g. insulin release in pancreatic beta cells (6) although a role for modulating the ER’s Ca^2+^ handing was not explored in those studies.

To date, the most well-known ER-PM junction with respect to Ca^2+^ handling is formed between the ER-lumen Ca^2+^ sensor STIM1 and the Ca^2+^ channel Orai, which are the fundamental working machinery of the CRAC channel. In the classical pathway for store-operated calcium entry (SOCE), the CRAC channel is formed only when the ER lumen Ca^2+^ concentration is dramatically depleted and transiently exists until the ER lumen is filled. Kv2.1 could also enhance this process as suggested by the somatic ER refilling data of Fig. S3. However, in the context of neuronal signaling, the process of activating a CRAC channel is rather slow and can cause Ca^2+^ spillover into the cytosol when activated; it is difficult to imagine neurons relying on this mechanism alone to maintain ER Ca^2+^. Indeed, chronic activation of CRAC channels was found to upregulate spontaneous vesicle fusion (54). Additionally, activation of STIM appears to inhibit or activate a number of PM proteins (55). Thus, although STIM1 and Orai can replenish ER stores when the ER lumen is severely depleted of Ca^2+^, an “on-demand” mechanism to efficiently keep the ER filled and coupled to electrical activity solves several problems without perturbing additional novel sources of Ca^2+^ entry and spillover. In this way, the Kv2.1-VAP ER-PM junctions are different than CRAC channels in that they are engaged independent of ER-lumen Ca^2+^ levels. Interestingly, Kv2.1 clusters are dynamic in some situations and can be dispersed in hypoxic conditions such as a stroke (56, 57) or when exposed to high levels of extracellular glutamate (12, 56, 58). This may be quite useful for limiting neurotransmitter release under certain pathological conditions that will require additional experiments.

Why form ER-PM junctions with a voltage-sensitive protein? The movement of the positive charges within the voltage sensor during membrane depolarization produces a gating current that precedes and is independent of ion conduction during channel opening. It is possible that electrically excitable cells could be using Kv2.1 as a voltage sensor to communicate changes in membrane potential across the ER-PM junction just as the charge movement of the voltage sensor in L-type Ca^2+^ channels communicates to ryanodine receptors in mammalian skeletal muscle during excitation-contraction coupling (59, 60). Although we cannot fully rule out communication of charge movement from Kv2.1 to the ER initiating Ca^2+^ uptake, it seems unlikely as our use of the gating modifier GxTx, which should block both voltage-sensing and conduction, did not impair neurotransmission. It has also been shown that ion channels have preferred lipid environments, so the localization of ER-PM junctions by an ion channel could also help direct the junctions to favorable areas within the heterogeneous lipid environment of the PM.

As the Kv2.1-VAP mediated ER-PM junction allows the ER to quickly access Ca^2+^ during electrical activity, a key question remains as to what the ER is doing with the additional Ca^2+^. It has been proposed that an essential role of the ER is to shuttle Ca^2+^ to other organelles, including the mitochondria Ca^2+^ uniporter (MCU), which has different affinities for Ca^2+^ depending on subunit expression. A recently identified MICU3 subunit is required for neuronal mitochondria to receive Ca^2+^ from the cytosol (61), but it remains unclear if they also receive Ca^2+^ from the ER to couple electrical activity to ATP production. A second possibility is that the ER may be shuttling Ca^2+^ to other organelles, or Ca^2+^-sensitive proteins. Indeed, the cytosol of the neuron is one of the most highly buffered Ca^2+^ environments identified between cells; thus, a mechanism to coordinate delivery to microdomains of Ca^2+^ within the cytoplasm may be very useful for coupling Ca^2+^ signaling to protein activation in distal processes of neurons like the axon. One example is seen in *Drosophila*, where ER Ca^2+^ is used to activate calcineurin, a Ca^2+^-dependent phosphatase, in an essential role for synapse development (62). Alternatively, rather than acting to move Ca^2+^ between organelles, the ER may be polarized within the neuron taking in Ca^2+^ during electrical stimulation, which may then be released at other subcellular locations within the soma or axon. As the dynamics of ER Ca^2+^ are not static, our identification of a novel, on demand mechanism for ER Ca^2+^ filling during electrical stimulation opens new avenues of research to understand ER signaling in neurons.

## Acknowledgements

We thank Samuel Bergerson, Michelle Gleason, and Amelia Ralowicz for critical reading of the manuscript. This work was supported by the Esther A. and Joseph Klingenstein Fund (M.B.H.), NIH NINDS grant F31NS110192 (L.C.P.), NIH NINDS grant 1R01NS112365 (M.M.T. and M.B.H.), and P20 NIGMS grant GM113132 (M.B.H.).

## Materials and Methods

### Cell Culture and Transfection

Primary neurons from postnatal day 0-1 Sprague Dawley rats of either sex were cultured for all experiments. Briefly, hippocampal CA1-CA3 regions with the dentate gyri removed were harvested, tissue was dissociated into single cells with bovine pancreas trypsin and cells were plated onto poly-L-ornithine-coated glass coverslips inside a 6mm cloning cylinder. Ca^2+^ phosphate-mediated DNA transfection was performed on cultures at 5-6 days *in vitro* (DIV). In some cases, GFP-Kv2.1 and dsRedER were transfected using Lipofectamine 2000 (Life Technologies) as previously described (12). All experiments were performed on mature neurons between 14-24 days *in vitro* unless noted otherwise. To ensure reproducibility, experiments were performed on neurons from a minimum of three separate cultures. All protocols used were approved by the Institutional Animal Care and Use Committee at Dartmouth College and conform to the NIH Guidelines for the Care and Use of Animals.

### Genetic Tools

The following constructs were used: QuasAr2 (variant DRH334, hSyn promoter) (28), vGlut-pHluorin (43), SypGCaMP6f (48, 49), ER-GCaMP6-150 (Addgene #86918) (20), CD4-Kv2.1FFAT (11), mGreenLantern (Addgene #161912) (33), and R-CEPIA1er (Addgene #58216) (38). To reduce endogenous Kv2.1 expression for live cell imaging, an shRNA plasmid was obtained from OriGene against the following mRNA target sequence: CAGAGTCCTCCATCTACACCACAGCAAGT. For rescue experiments in Kv2.1 knockdown neurons, three silent mutations were introduced into the rat Kv2.1 sequence to generate the “wobbled” Kv2.1 (wKv2.1) construct: C2154T, C2157A, C2160T.

### Live Cell Imaging

All experiments were performed at 34°C using a custom-built objective heater. Cultured cells were mounted in a rapid-switching laminar flow perfusion and stimulation chamber on the stage of a custom-built epifluorescence microscope. Neurons were continuously perfused at a rate of 400µl/min in a modified Tyrode’s solution containing the following (in mM): 119 NaCl, 2.5 KCl, 2 CaCl_2_, 2 MgCl_2_, 25 HEPES, and 30 glucose with 10 μM CNQX (Sigma-Aldrich) and 50 μM AP5 (Sigma-Aldrich). Images were obtained using either a Zeiss Observer Z1 equipped with an EC Plan-Neofluar 40x 1.3 NA oil immersion objective, or an Olympus IX-83 microscope equipped with a 40x 1.35 NA oil immersion objective (UApoN40XO340-2). All images were captured with an IXON Ultra 897 EMCCD (Andor) that was cooled to -80°C by an external liquid cooling system (EXOS). All excitation light occurred via OBIS lasers (Coherent). APs were evoked by passing 1 ms current pulses yielding fields of ∼12 V/cm^2^ via platinum/iridium electrodes. Timing of stimulation was delivered by counting frame numbers from a direct readout of the EMCCD rather than time itself for more exact synchronization using a custom-built board powered by an Arduino Duo chip manufactured by an engineering firm (Sensorstar).

#### Voltage Measurements

QuasAr fluorescence was recorded with a 980 µs exposure time; images were acquired at 1 kHz using an OptoMask (Cairn Research) to prevent light exposure of non-relevant pixels. Cells were illuminated with 70 – 120 mW by an OBIS 637 nm laser (Coherent) with ZET635/20×, ET655lpm, and ZT640rdc filters (Chroma). For somatic recordings, 25 AP stimulations delivered a 4 Hz were trial averaged, and for axonal recordings 100 AP stimulations delivered at 4 Hz were trial averaged.

#### Cytosolic Calcium Measurements

GCaMP6f fluorescence was recorded with a 29.5 ms exposure time and images were acquired at 30 Hz. Cells were illuminated by an OBIS 488 nm laser at 7-9 mW (Coherent) with ET470/40x, ET525/50 m, and T495lpxr filters (Chroma). We repeated ad averaged 3-4 trials to measure AP train stimulation-induced responses.

#### ER Calcium Measurements

ER-GCaMP6-150 fluorescence was recorded with a 19.8 ms exposure time and images were acquired at 50 Hz. Cells were illuminated by an OBIS 488 nm laser at 7-9 mW for axonal recordings, 1-2 mW for somatic recordings (Coherent) with ET470/40x, ET525/50 m, and T495lpxr filters (Chroma). We repeated and averaged 3-4 trials measure AP train stimulation-induced responses.

#### Vesicle Fusion Measurements

vGlut-pHluorin fluorescence was captured with an exposure time of 9.8 ms and images were acquired at 100 Hz. Cells were illuminated by an OBIS 488 nm laser at 7-9 mW (Coherent) with ET470/40x, ET525/50 m, and T495lpxr filters (Chroma). For Guangxitoxin experiments, GxTx was continuously perfused at 100 nm (Alomone Labs). Cells were bathed in 50 mM NH_4_Cl to neutralize vesicle pH at the end of each experiment to quantify vesicle pool size.

### Immunocytochemistry

#### To validate the effectiveness of our knockdown strategy

Neurons were fixed with 4% paraformaldehyde and 4% sucrose in phosphate-buffered saline (PBS) for 10 minutes, permeabilized with 0.2% Triton X-100 for 10 minutes and blocked with 5% goat serum/5% bovine serum albumin in PBS for 30 minutes at room temperature. Neurons were then incubated with the Kv2.1 primary antibody K89/34 (1:500, NeuroMab) and the GFP primary antibody A10262 (1:1000, Invitrogen) overnight at 4°C. Cells were washed three times with PBS and incubated 1 hour at room temperature with Alexa Fluor-conjugated secondary antibodies (1:1000, Invitrogen). All antibodies were diluted in 5% goat serum for incubation.

#### To detect the localization of endogenous Kv2 channels

Hippocampal cultures of neurons were isolated from E18 Sprague Dawley rat brains of both sexes. Pregnant rats were deeply anaesthetized with isoflurane, as outlined in a protocol approved by the Institutional Animal Care and Use Committee of Colorado State University (protocol ID: 15-6130A). Hippocampi were dissociated and cultured as previously described for neurons (63, 64). Cultures were seeded on glass-bottom 35mm dishes with No. 1.5 coverslips (MatTek, Ashland, MA) coated with poly-L lysine (Sigma-Aldrich, St. Louis, MO) in borate buffer, and in a medium composed of Neurobasal (Gibco/Thermo Fisher Scientific, Waltham, MA), B27 Plus Supplement (Gibco/Thermo Fisher Scientific, Waltham, MA), Penicillin/Streptomycin (Cellgro/Mediatech, Manassas, VA), and GlutaMAX (Gibco/Thermo Fisher Scientific).

Cultures of the indicated day *in vitro* (DIV) were fixed with 4% formaldehyde for 15 min at room temperature in neuronal imaging saline (NIS) composed of 126 mM NaCl, 4.7 mM KCl, 2.5 mM CaCl_2_, 0.6 mM MgSO_4_, 0.15 mM NaH_2_PO_4_, 0.1 mM ascorbic acid, 8 mM glucose, and 20 mM HEPES, pH 7.4, 300 mOsm. Following six washes with NIS the fixed cells were blocked in NIS with 10% goat serum and 0.1% Triton X-100 for 4-10 hrs at room temperature. Purified Kv2 mouse monoclonal antibodies, knockout verified, were from NeuroMab (Davis, CA (Kv2.1, K89/34 and Kv2.2, N37B/1) and used at a 1/1000 dilution in NIS with 10% goat serum and 0.1% Triton X-100 for 1 hr at room temperature followed by three 5 min washes in NIS and then a 45 min secondary antibody (1/2000 dilution) incubation at room temperature in NIS with 10% goat serum and 0.1% Triton X-100. The Alexa Fluor 488-conjugated goat anti-mouse IgG (A11001) secondary antibody was from Invitrogen (Walthan, MA). Cells were then rinsed three times for 5 min each, immediately mounted under glass coverslips with Aqua-Poly/Mount (Polysciences, Warrington, PA), and imaged as described below. Synapsin 1/2 rabbit polyclonal antibody (002 106) was obtained from Synaptic Systems (Goettingten, Germany) and used at 1/2000 as described above except that here the secondary antibody was Alexa Fluor 647-conjugated goat anti-rabbit IgG (A21244), also from Invitrogen.

Spinning disk confocal microscopy was performed on immunolabeled cultures using a Yokogawa (Musashino, JP) based CSUX1 system with an Olympus (Tokyo, JP) IX83 inverted stand, and coupled to an Andor (Abingdon, GB) laser launch containing 405, 488, 568, and 637 nm diode lasers, 100-150 mW each. Images were collected using an Andor iXon EMCCD camera (DU-897) and 100X Plan Apo, 1.4 NA objective. To prevent bleed over between fluorescent channels all imaging was performed sequentially with paired excitation-emission filter settings. This confocal system uses MetaMorph software (version 7.8.13.0). Image analysis and presentation was performed using Volocity v6.1.1 software and all images were filtered and adjusted for brightness and contrast.

### Western Blotting

To verify the efficiency of Kv2.1 knockdown, hippocampal neurons were treated at DIV 3 with AAV1mCherry-pU6-Kv2.1 shRNA designed by OriGene (target sequence: CAGAGTCCTCCATCTACACCACAGCAAGT) and produced by Vector Biolabs. Untreated neurons were used as a control. At DIV 19, control and Kv2.1 shRNA virus-treated neurons were extracted in radioimmunoprecipitation assay (RIPA) buffer (50mM Tris, pH 8.0, 150mM NaCl, 1% Nonidet P-40 (NP-40), 0.5% sodium deoxycholate, and 0.1% SDS supplemented with protease inhibitors) for 1 hour at 4°C. The lysates were centrifuged at 13000g at 4°C for 10 min, and the protein concentrations were measured with bicinchoninic acid assays. The proteins were loaded and run on SDS-PAGE gels, transferred to PVDF membranes, and immunoblotted with anti-Kv2.1 antibody (K89/34, NeuroMab) and anti-α tubulin antibody (Sigma). The blots were detected using chemiluminescence reagent on a Bio-Rad Chemidoc MP and Image J was used for quantification of detected bands.

### Image and Data Analysis

Images were analyzed in ImageJ using a custom-written plugin (http://rsb.info.nih.gov/ij/plugins/time-series.html). To quantify fluorescence, we selected 1.4µm diameter circular regions of interest (ROIs) from ΔF images of each experiment, recentering on the brightest pixel within the ROI in the ΔF image. ROIs were selected based on localized responses of voltage, calcium, or vesicle fusion, rather than morphology, to define a presynaptic terminal. All statistical data are presented as means +/-SEM (n = number of neurons) and all experiments were performed on more than three independent cultures. To measure the full-width at half maximum of QuasAr fluorescence, we used Origin version 9.1 (Origin Lab). Quantification of vesicle fusion was obtained by normalizing the fluorescence change in response to stimulation to the total number of vesicles measured by application of ammonium chloride.

### Quantification and Statistical Analysis

Statistical analyses were performed in Excel and Origin. We used paired two sample for means *t*-test for paired results. Normally distributed data were processed with the Student’s *t*-test for two independent distributions.

---

**Supplementary Figure 1.**
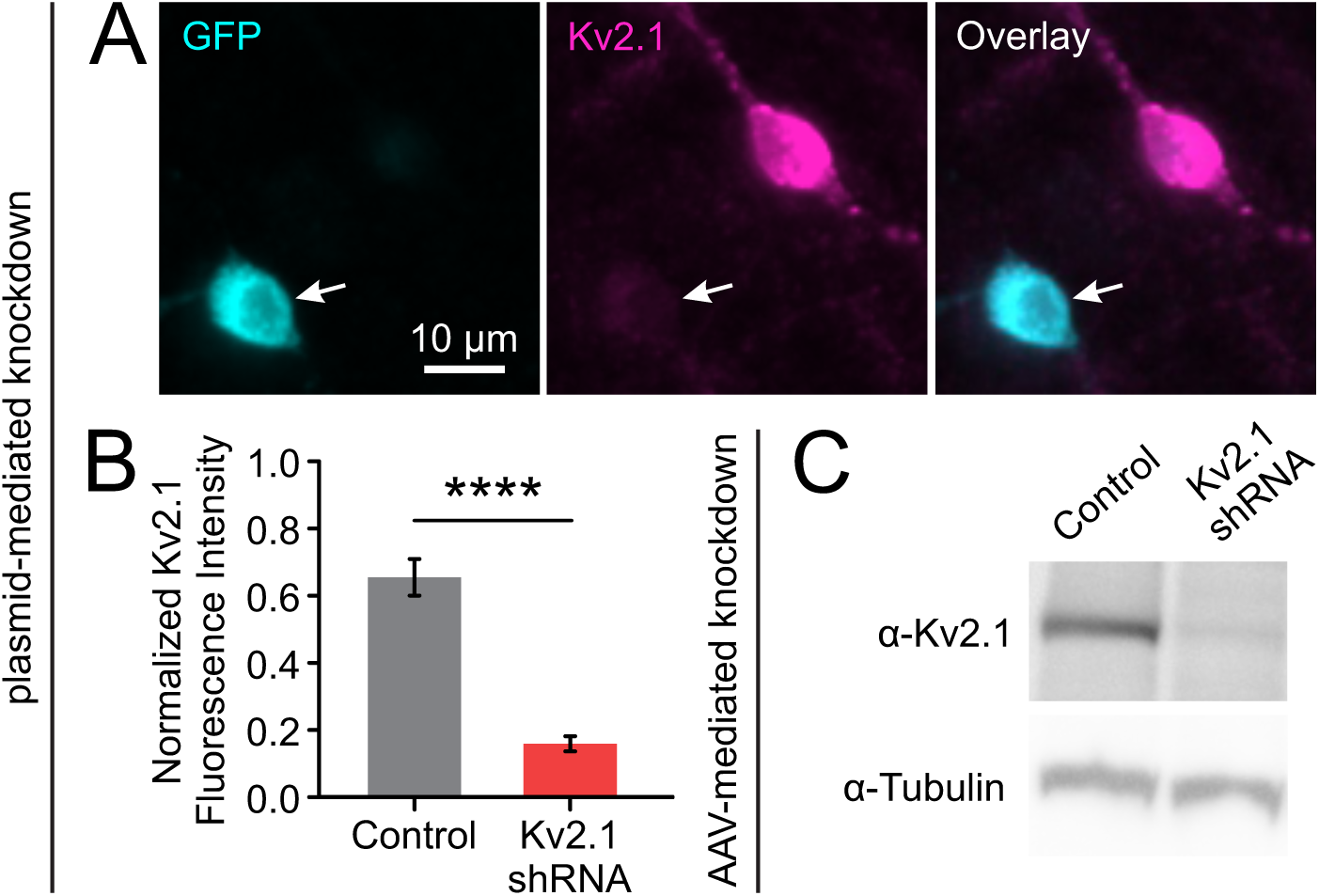
Kv2.1 expression is reduced with genetic knockdown. Kv2.1 shRNA was designed to efficiently remove not only Kv2.1 conductance but the protein itself to examine both conducting and non-conducting roles. Plasmid encoding shRNA was delivered using Ca^2+^ phosphate-based transfection to a small fraction of neurons in culture (<1% of total neurons). This allows comparison of protein expression levels of each cell with adjacent untransfected neurons to examine the efficiency of protein knockdown. **(A-B)** Cultured hippocampal neurons were immunostained for GFP and endogenous Kv2.1.Arrows mark a neuron co-transfected with GFP and Kv2.1 shRNA that is depleted of endogenous Kv2.1. **(B)** Quantification of immunostained fluorescence intensity (Control neurons, *n* = 20 cells; Kv2.1 KD neurons, *n* = 38 neurons; *p* < 0.0001, Student’s *t*-test). We also developed an adeno-associated virus (AAV) to deliver this same shRNA to quantify knockdown of total protein levels using a Western blot **(C)** for neurons with AAV-mediated Kv2.1 knockdown by shRNA with α-tubulin as a loading control to mark total protein content.

**Supplementary Figure 2.**
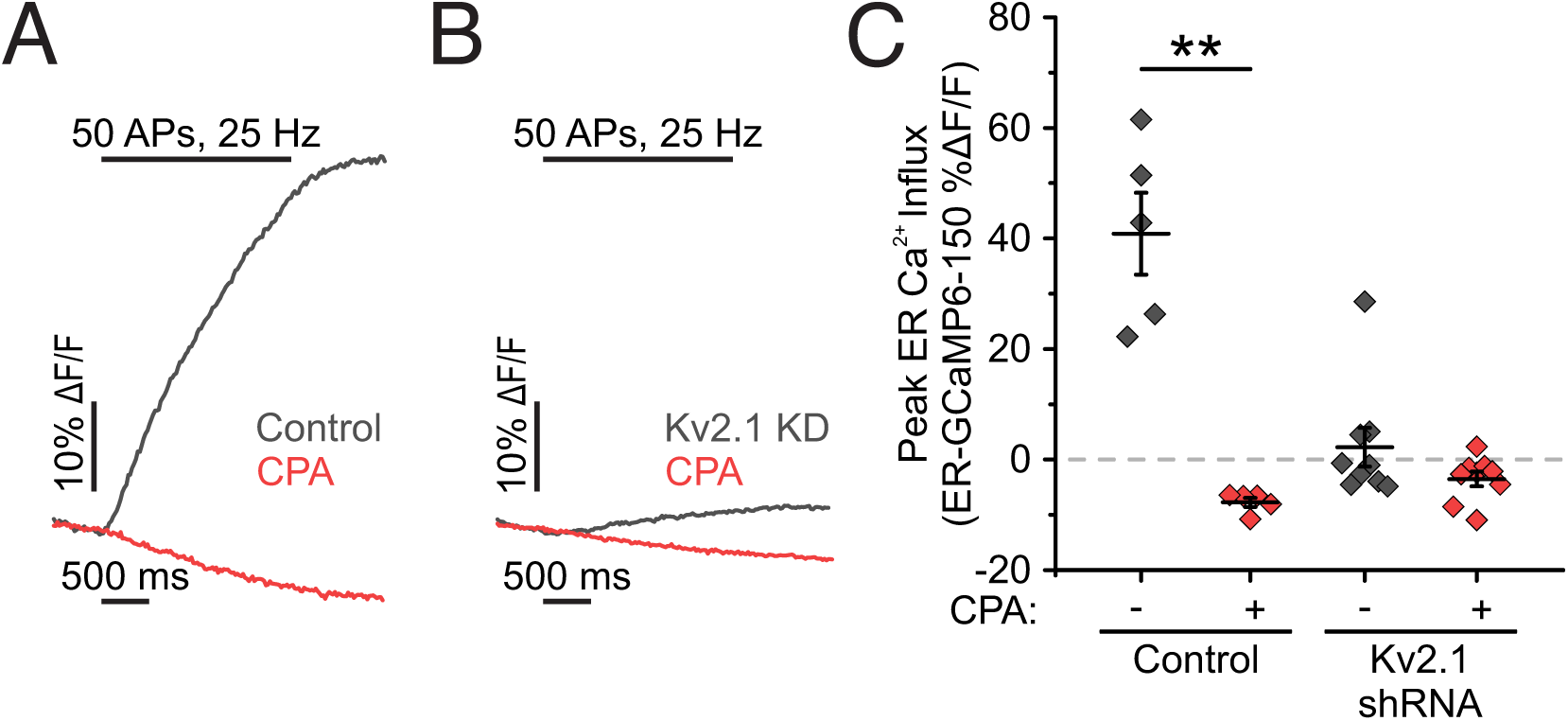
ER Ca^2+^ uptake is mediated by SERCA pumps. The only known major conduit for Ca^2+^ to enter the ER is through the SERCA pumps. As such, we used the specific inhibitor cyclopiazonic acid (CPA) to confirm that Kv2.1 was enabling uptake of cytosolic Ca^2+^ through SERCA pumps. **(A-B)** Average fluorescence traces of somatic ER-GCaMP6f-150 with CPA treatment in control **(A)** and Kv2.1 knockdown **(B)** neurons. **(C)** Quantification of peak fluorescence (Control neurons, *n* = 5 cells; Kv2.1 KD neurons, *n* = 9 cells; *p* < 0.01, paired *t*-test).

**Supplementary Figure 3.**
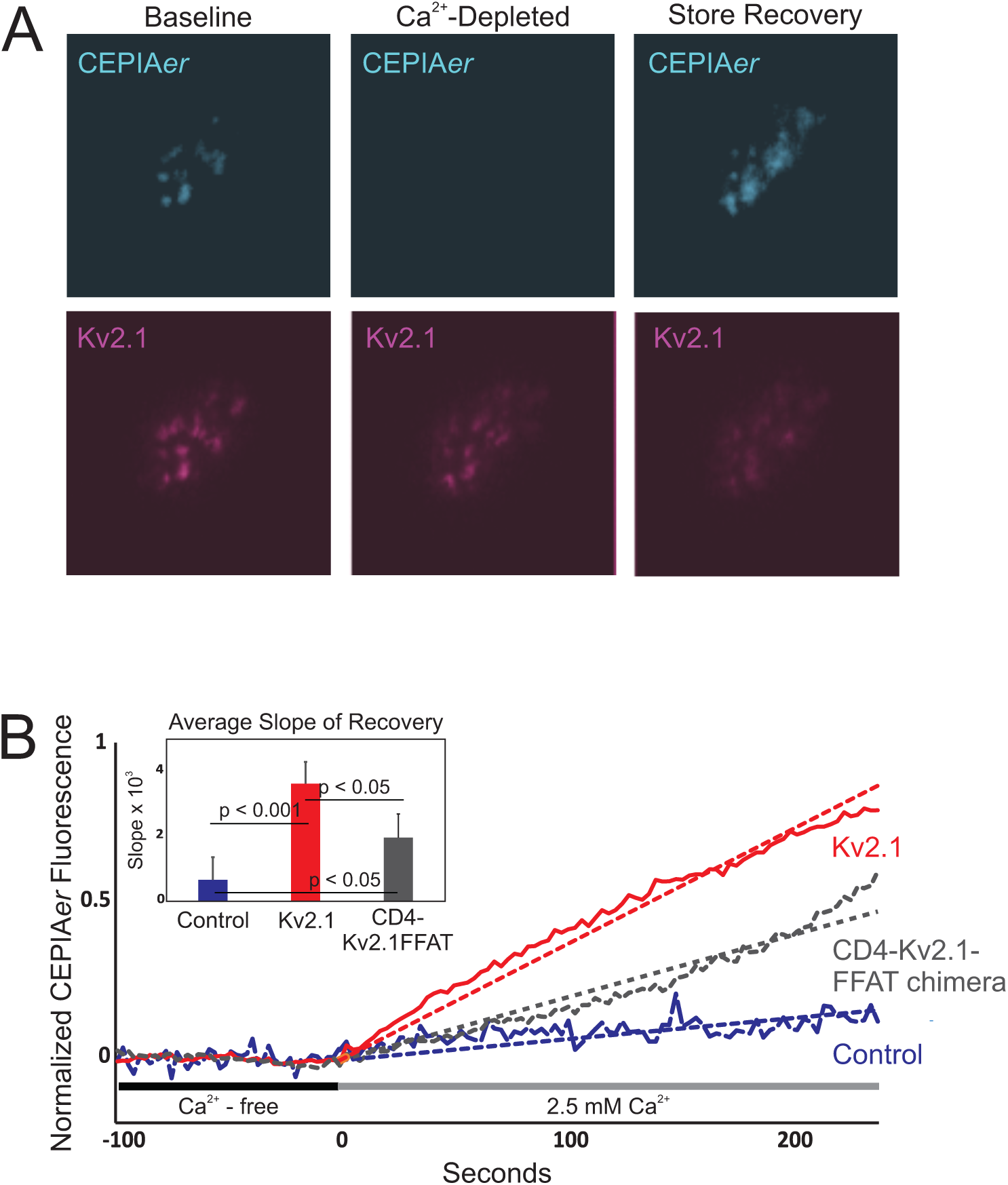
Somatic ER Ca^2+^ refilling following depletion is enhanced in the presence of Kv2.1. The inability of the ER to take up Ca^2+^ during AP stimulation revealed a specific role for Kv2.1 to couple Ca^2+^ handling with electrical stimulation. We also tested a potential role for Kv2.1 in facilitating Ca^2+^ uptake into the ER after depletion of Ca^2+^ from the ER by a prolonged removal of extracellular Ca^2+^. Young DIV 7 neurons have two advantages to deploy in this study: first, they have very low endogenous expression of Kv2.1; second, they are still growing on the glass coverslip prior to astrocyte expansion and interpolation beneath the neurons. As such they are amenable to high-resolution imaging of ER-PM junctions using total-internal reflection microscopy (TIRF). This method revealed that full-length Kv2.1 generates efficient ER-calcium refilling, while simply expressing the CD4-Kv2.1FFAT motif chimera, which forms ER/PM junctions and concentrates VAPs here without other Kv2.1 sequence (11), only partially enhances ER Ca^2+^ refilling, perhaps due to simply increasing ER/PM contact area. **(A)** Examples of TIRF-based ER Ca^2+^ imaging with CEPIA*er* at Kv2.1-induced ER/PM contact sites before, immediately after store depletion, and after store recovery. **(B)** Refilling rates at different ER/PM junctions, averaged from four independent measurements per condition.

**Supplementary Figure 4.**
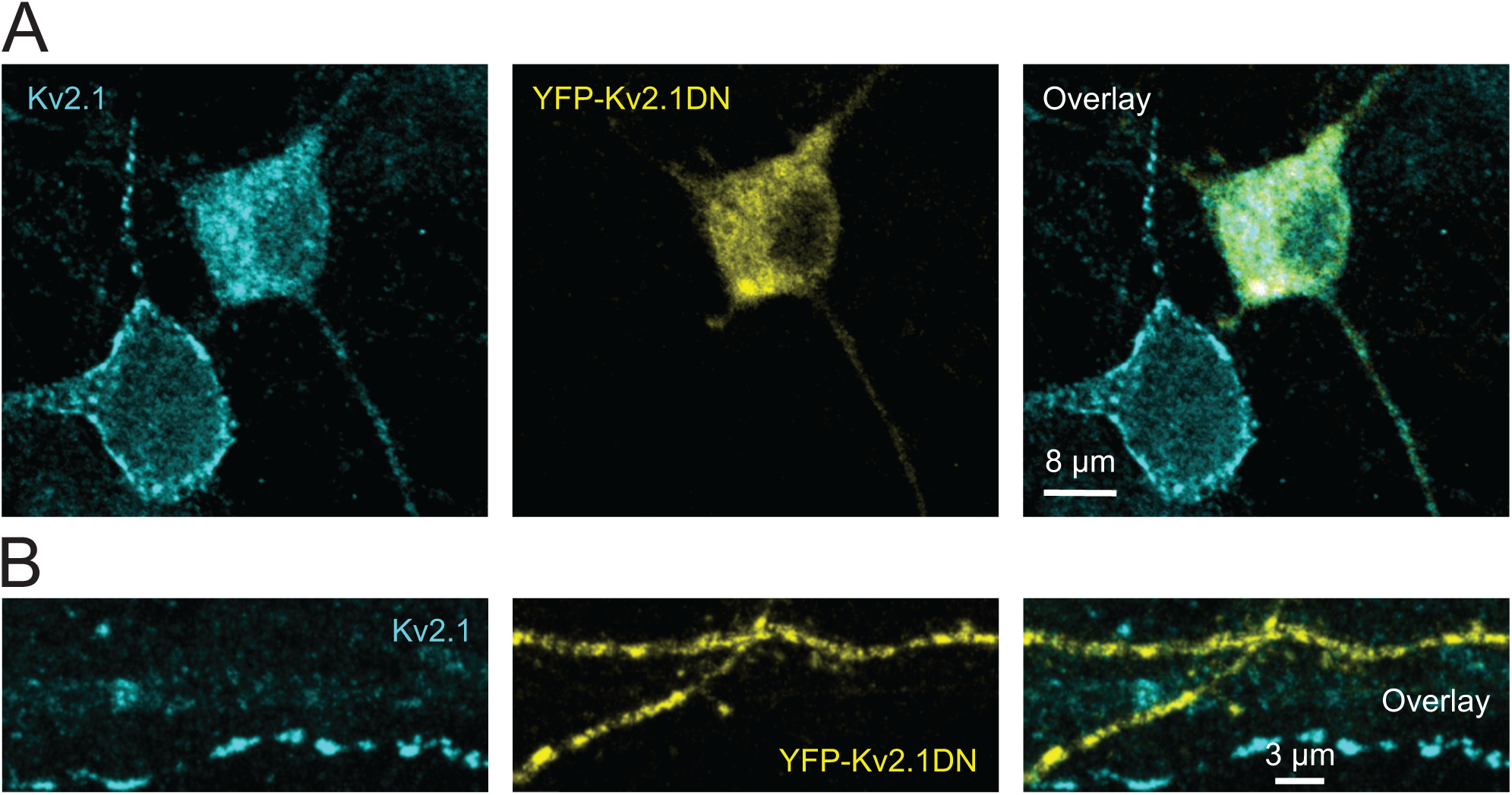
Dominant-negative verification of endogenous Kv2.1 immunolocalization to axons. Expression of a YFP-tagged dominant-negative form of Kv2.1 (YFP-Kv2.1DN, composed of just the YFP-tagged Kv2.1 N-terminus and first transmembrane domain) prevents trafficking of Kv2.1 tetramers to the neuronal surface. We used this construct to validate the immunolabeling of endogenous Kv2.1 by comparing YFP-Kv2.1DN-transfected neurons with untransfected neighbors. **(A)** Examples of immunolabeled endogenous Kv2.1 (*cyan*) and YFP-Kv2.1 DN (*yellow*) in the somas of two DIV 16 neurons. Note the membrane expression of Kv2.1 in the neuron on the left without YFP, while the neuron expressing YFP-Kv2.1DN shows Kv2.1 restricted to the interior of the cell body where the DN-endogenous Kv2.1 complex likely remains trapped in the ER. **(B)** Examples of immunolabeled endogenous Kv2.1 and YFP-Kv2.1DN in two adjacent axons.

**Supplementary Figure 5.**
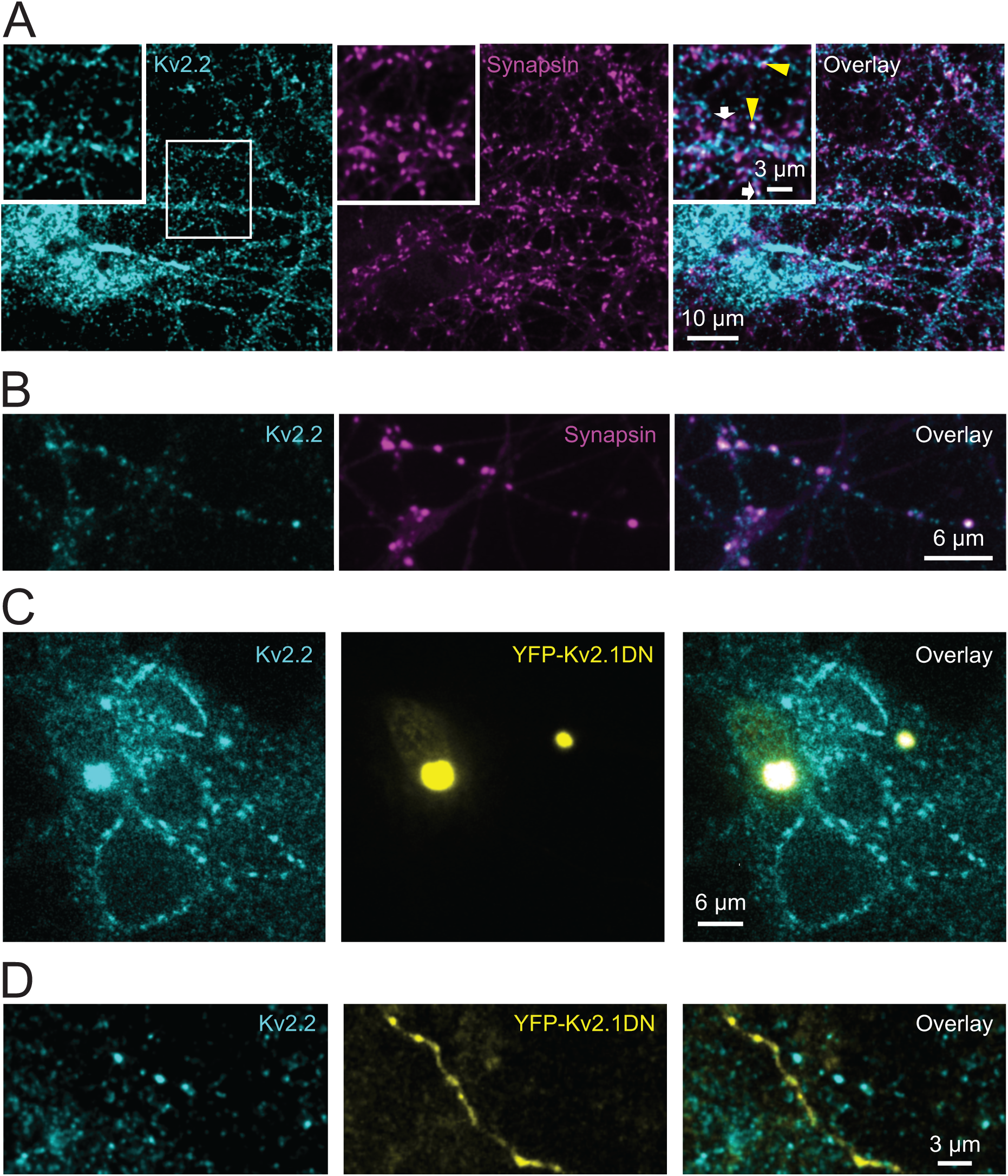
Immunolocalization of endogenous Kv2.2. **(A)** Immunolabeling of endogenous Kv2.2 (*cyan*) and synapsin (*magenta*), with merged channels (*right*), in DIV 20 cultured hippocampal neurons. The center white box indicates the region enlarged as shown in the inset. The white arrows indicate Kv2.2 colocalized with synapsin-positive presynaptic terminals and the yellow arrowheads point to areas where the colocalization is less exact. **(B)** Colocalization of endogenous Kv2.2 with synapsin in an isolated axon. Expression of the dominant-negative form of Kv2.1 (YFP-Kv2.1DN), which also assembles with Kv2.2, was again used to validate the immunolabeling of endogenous Kv2.2 by comparing DN-transfected with untransfected neurons. **(C-D)** Examples of immunolabeled endogenous Kv2.2 (*cyan*) in neurons with and without the expressed YFP-Kv2.1DN (*yellow*) in somas and axons, respectively, of neurons at DIV 14. In **(C)**, note the cell surface expression of Kv2.2 in the YFP-Kv2.1 DN-free neurons and the dramatic accumulation of Kv2.2 with the DN in the interior of the two YFP-positive cells.

**Supplementary Figure 6.**
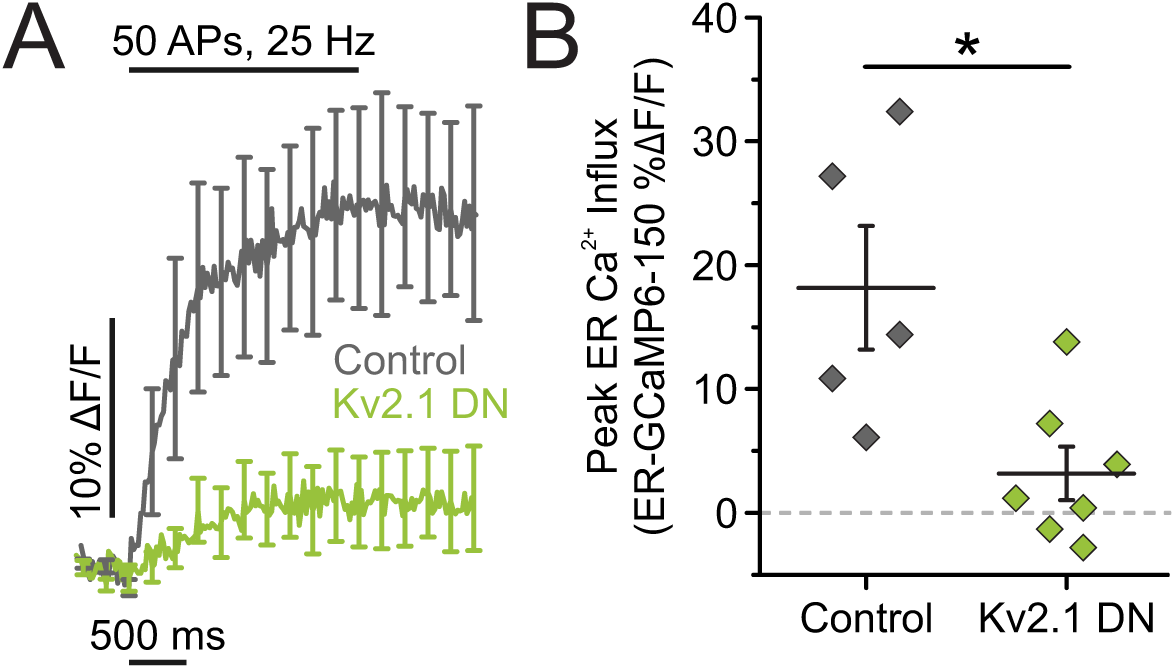
Genetic knockdown with a Kv2.1 dominant-negative subunit decreases ER Ca^2+^ influx. To determine what contribution Kv2.2 could potentially make to enable ER Ca^2+^ uptake in axons, we expressed the dominant negative form of Kv2.1 to remove both Kv2.1 and Kv2.2 from the membrane surface. This impaired ER Ca^2+^ uptake to the same extent as removing Kv2.1 alone, suggesting a minor or non-existent role for Kv2.2 in modulating stimulation-evoked ER Ca^2+^ uptake in hippocampal axons. **(A-B)** Average fluorescence traces of axonal ER-GCaMP6f-150 **(A)** and quantification of peak fluorescence **(B)** for both control and Kv2.1 DN neurons (Control neurons, *n* = 5 cells; Kv2.1 DN neurons, *n* = 7 cells; *p*<0.05, Student’s *t*-test).

**Supplementary Figure 7.**
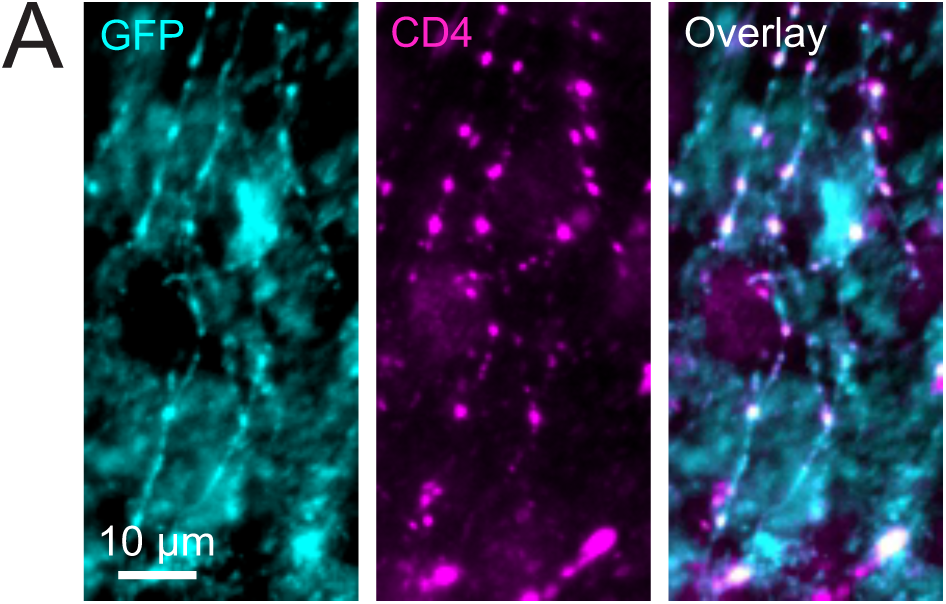
CD4-Kv2.1FFAT traffics to the distal axon. Although CD4 protein trafficking is not typically subject to polarized trafficking, we wanted to make sure that CD4-Kv2.1FFAT expression patterns were still robust in the axons. Indeed, punctate fluorescence was visible in neurons simply expressing the chimera to confirm axonal trafficking. **(A)** Neurons expressing Kv2.1 shRNA and CD4-Kv2.1FFAT were immunolabeled against CD4 and GFP as a transfection marker. Scale bar 10μm.

